# Integrin Affinity Modulation Critically Regulates Atherogenic Endothelial Activation *in vitro* and *in vivo*

**DOI:** 10.1101/2020.02.14.949941

**Authors:** Zaki Al-Yafeai, Jonette M. Peretik, Brenna H. Pearson, Umesh Bhattarai, Dongdong Wang, Brian G. Petrich, A. Wayne Orr

**Affiliations:** Department of Molecular and Cellular Physiology, LSU Health Sciences Center, Shreveport, LA; Department of Cell Biology and Anatomy, LSU Health Sciences Center, Shreveport, LA; Department of Pathology and Translational Pathobiology, LSU Health Sciences Center, Shreveport, LA; Department of Pediatrics, Emory University, Atlanta, GA

**Author notes:** Corresponding author: A. Wayne Orr, Department of Pathology and Translational Pathobiology, 1501 Kings Hwy, Biomedical Research Institute, Rm. 6-21, LSU Health Sciences Center – Shreveport, Shreveport, LA 71130.

## Abstract

While vital to platelet and leukocyte adhesion, the role of integrin affinity modulation in adherent cells remains controversial. In endothelial cells, atheroprone hemodynamics and oxidized lipoproteins drive an increase in the high affinity conformation of α5β1 integrins in endothelial cells *in vitro*, and α5β1 integrin inhibitors reduce proinflammatory endothelial activation to these stimuli *in vitro* and *in vivo*. However, the importance of α5β1 integrin affinity modulation to endothelial phenotype remains unknown. We now show that endothelial cells (talin1 L325R) unable to induce high affinity integrins initially adhere and spread, but show significant defects in nascent adhesion formation. In contrast, overall focal adhesion number, area, and composition in stably adherent cells are similar between talin1 wildtype and talin1 L325R endothelial cells. However, talin1 L325R endothelial cells fail to induce high affinity α5β1 integrins, fibronectin deposition, and proinflammatory responses to atheroprone hemodynamics and oxidized lipoproteins. Inducing the high affinity conformation of α5β1 integrins in talin1 L325R cells partially restores fibronectin deposition, whereas NF-κB activation and maximal fibronectin deposition require both integrin activation and other integrin-independent signaling. In endothelial-specific talin1 L325R mice, atheroprone hemodynamics fail to promote inflammation and macrophage recruitment, demonstrating a vital role for integrin activation in regulating endothelial phenotype.

## Introduction

In the arterial microenvironment at atherosclerosis prone regions, local hemodynamics and matrix composition critically regulate endothelial activation and early atherosclerosis(1, 2). Disturbed flow primes endothelial cells to activation by systemic factors, in part through NF-κB activation(3, 4). However, the presence of a fibronectin-rich matrix significantly enhances disturbed flow-induced endothelial activation, whereas basement membrane proteins (collagen IV, laminin) support minimal endothelial activation(5–7). Similarly, matrix content plays a vital role in oxidized LDL (oxLDL)-mediated proinflammatory responses, where a fibronectin-rich matrix enhances endothelial activation in response to oxLDL(8, 9). Fibronectin support atherogenic endothelial activation through specific integrin signaling responses. Integrins, the largest family of receptors for extracellular matrix (ECM) proteins, are heterodimers of α and β subunits that mediate both cell-dependent extracellular matrix remodeling and matrix-dependent changes in cell phenotype(10). Blocking fibronectin-binding integrins suppresses endothelial activation by oxLDL and by disturbed flow both *in vitro* and *in vivo(8, 9, 11, 12)*, suggesting that cell-matrix interactions serve as an essential component of early atherogenic endothelial activation.

Integrins exist in multiple activation states, including a bent, closed state (low affinity), an intermediate extended state with a closed headpiece (intermediate affinity), and a fully extended, open conformation (high affinity)(13, 14). The intermediate affinity of extended, closed integrins mediate weak interactions, such as during integrin-mediated leukocyte slow rolling(15), whereas the extended, open conformation is required for high affinity interactions, such as leukocyte firm adhesion. The common final step in integrin affinity modulation involves interactions between the integrin β subunits cytoplasmic tail and talin1(16), a cytoskeletal adaptor protein with a head domain that binds the integrin β tail and a rod domain that binds the actin cytoskeleton(17). The talin1 head domain binds to an NPxY motif on the β integrin subunit driving a transition from the bent, closed to the extended, closed (intermediate affinity) conformation. A second interaction between the talin1 head domain and the β tail membrane-proximal region (MPR) drives the final transition to an extended, open (high affinity) conformation(18). A talin1 mutant (W359A) that prevents talin1 binding to the β integrin NPxY motif prevents integrin extension and integrin–mediated slow rolling(19, 20). A mutation in the talin1 head domain that prevents talin1 interactions with the integrin β subunit MPR (L325R) allows talin1 interaction with integrins and actin but prevents the transition to the extended, open (high affinity) conformation(19, 21, 22). Mice expressing only talin1 L325R in platelets and neutrophils show defects in thrombus formation and neutrophil firm adhesion, respectively, although integrin-mediated neutrophil slow rolling remains intact(19, 21).

Endothelial integrins mediate distinct responses based on integrin-ligand pair involved(23). The fibronectin-binding integrin α5β1 plays a clear role in atherogenic endothelial activation. Atheroprone areas show elevated α5 mRNA and protein expression(24, 25), and both disturbed flow and oxLDL induce α5β1-dependent fibronectin matrix assembly and endothelial proinflammatory responses including NF-κB activation and ICAM-1/VCAM-1 expression(8, 9, 26). Mice deficient for endothelial α5 integrin signaling show reduced atherogenic endothelial activation and diminished plaque formation(8, 27). In unstimulated cells, a majority of α5β1 exists in the bent, closed (low affinity) conformation(14), and both oxLDL and flow enhance α5β1 affinity for ligand(8, 28). However, the role of talin1-dependent integrin affinity modulation in endothelial cell function remains poorly defined. Global knockout of talin1 is embryonic lethal at 8.5-9.5 days and associated with vascular defects(29), and inducible talin1 deletion in endothelial cells results in an unstable intestinal vasculature, resulting in hemorrhage and premature death(30). However, these effects may be due to talin1’s role in linking integrins to the cytoskeleton rather than integrin affinity modulation. Therefore, we sought to utilize endothelial cells expressing only talin1 L325R to characterize the importance of talin1-dependent integrin affinity modulation in endothelial phenotype in the context of atherogenic endothelial activation.

## Results

### Talin1 L325R mutant endothelial cells lack oxLDL and flow-induced α5β1 integrin affinity modulation

To characterize the role of integrin affinity modulation in endothelial cell function, we isolated mouse lung endothelial cells from Talin1^fl/L325R^ mice that express one floxed talin1 allele and one L325R mutant allele. Talin1^fl/L325R^ endothelial cells were then treated with adenovirus expressing either GFP (talin1 wildtype (WT)) or expressing Cre recombinase (talin1 L325R) to delete the floxed allele. While the talin1^Δ/L325R^ (hereafter talin1 L325R) endothelial cells contain only one copy of the talin1 gene, talin1^fl/L325R^ (hereafter talin1 WT) and talin1 L325R endothelial cells show equivalent protein levels of talin1 (**Figure 1A**). Consistent with previous reports in talin deficient cells(31), talin1 L325R endothelial cells show slowed adhesion and spreading compared to talin1 WT cells (**Figure 1B/C**). While talin1/2 deficient cells eventually lose the spread cell phenotype(31), talin1 L325R cells remain stably adherent, suggesting that the retained ability of talin1 L325R to link integrins to the cytoskeleton is sufficient to support stable cell adhesion. In contrast, talin1 L325R endothelial cells show significantly reduced formation of nascent adhesions at the leading edge of protrusions (**Figure 1D/E, Supplemental Figure Ia-c**), consistent with unstable protrusions and poor cell spreading. In isolated focal adhesion fractions, we observed that talin1 WT and talin1 L325R endothelial cells show similar recruitment of talin1, α5, and β1 integrins (**Figure 1F, Supplemental Figure IIa-e**), as well as the talin-binding proteins paxillin and vinculin (**Supplemental Figure III**). Immunocytochemistry for active β1 integrins (9EG7 antibody) and vinculin show that talin1 WT and talin1 L325R endothelial cells form similar number, size, and area of focal adhesions (**Figure 1G-J, Supplemental Figure IV**), suggesting that stable integrin adhesions are unaffected by preventing high affinity endothelial integrins.

**Figure 1.**
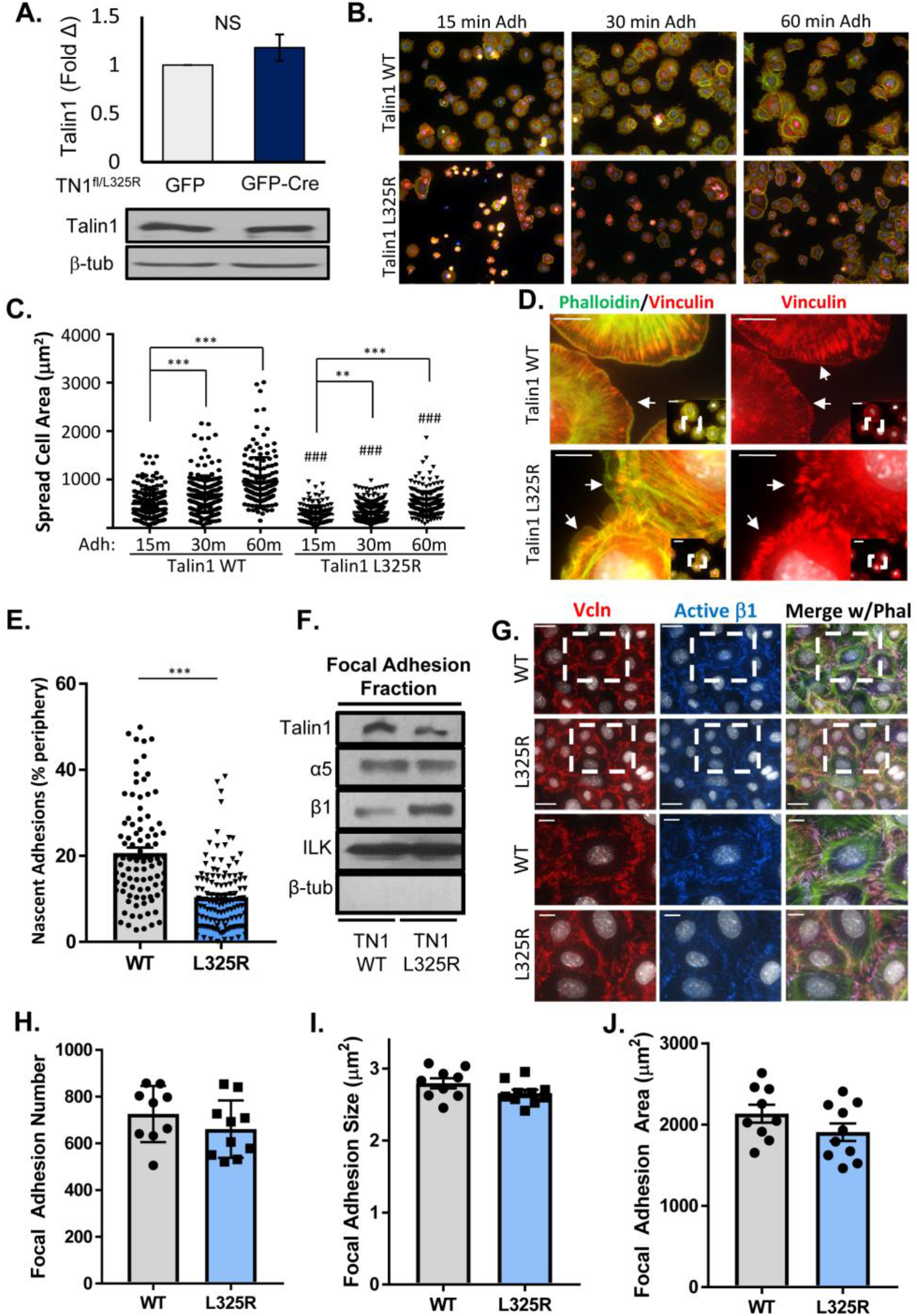
Talin1 L325R Endothelial Cells Show Deficient Spreading and Nascent Adhesions but Normal Focal Adhesion Structure. A) Representative Western blots of Mouse Lung Endothelial Cells (MLEC) from Talin1^fl/L325R^ after expression of GFP or GFP-Cre to delete the floxed allele. B-C) Talin1 WT and Talin1 L325R cells plated on fibronectin for 15, 30 and 60 minutes were fixed and immunostained for 9EG7 (red) and phalloidin 488 (green). Spread cell area was quantified for each cell across three independent experiments. Representative 20x micrographs are shown. D-E). Cells were stained as in (B) and nascent adhesions forming within 1 μm of the cell edge at 30 minutes adhesion were visualized and quantified. Representative micrographs are shown with 60x inset. Scale bar = 10 μm. F) Focal adhesion fraction in TN1 WT and TN1 L325R MLECs were blotted for different focal adhesion proteins. n=3. G-J) Talin1 WT and Talin1 L325R cells were plated on fibronectin for 2 hours and stained as in (B). Focal adhesion number, average size, and area per high powered field were quantified for each of three separate experiments. Representative micrographs are shown with 60x inset. Scale bar = 20 μm and 10 μm for image within boxed area. Values are means ±SE. Student t test was used for statistical analysis. ** p<0.01, *** p<0.001.

The presence of only the Talin1 L325R mutant impairs the induction of high affinity αIIbβ3 in platelets(21) and high affinity αLβ2 in leukocytes(19). Therefore, we assessed whether talin1 L325R endothelial cells show a similar reduction in α5β1 integrin affinity modulation in response to flow and oxLDL(8, 28). Talin1 WT and L325R endothelial cells were exposed to shear stress or oxLDL, and α5β1 affinity modulation was measured using the ligand mimetic GST-FNIII_9-11_, a GST fusion protein containing 9^th^ to 11^th^ type III repeats of fibronectin(28), that binds high affinity, unligated integrins. Retention of GST-FNIII_9-11_ is specifically mediated by α5β1 and is sensitive the both positive and negative regulators of integrin affinity modulation(28). While basal levels of GST-FNIII_9-11_ retention were similar between talin1 WT and talin1 L325R endothelial cells (**Figure 2A/B**), the enhanced levels of high affinity α5β1 observed in response to shear stress (**Figure 2A**), and oxLDL (**Figure 2B**) was absent in talin1 L325R endothelial cells. Immunostaining for active β1 (9EG7), which demonstrates both high affinity, unligated and high affinity, ligated integrins, showed enhanced active β1 staining in response to oxLDL and flow in talin1 WT but not talin1 L325R endothelial cells, consistent with a an impaired integrin affinity modulation response (**Figure 2C/D**). These results suggest that talin1 is dispensable for the formation of stable adhesions but is required for the induction of high affinity integrins in endothelial cells that contribute to nascent adhesion formation and the endothelial response to oxLDL and flow.

**Figure 2.**
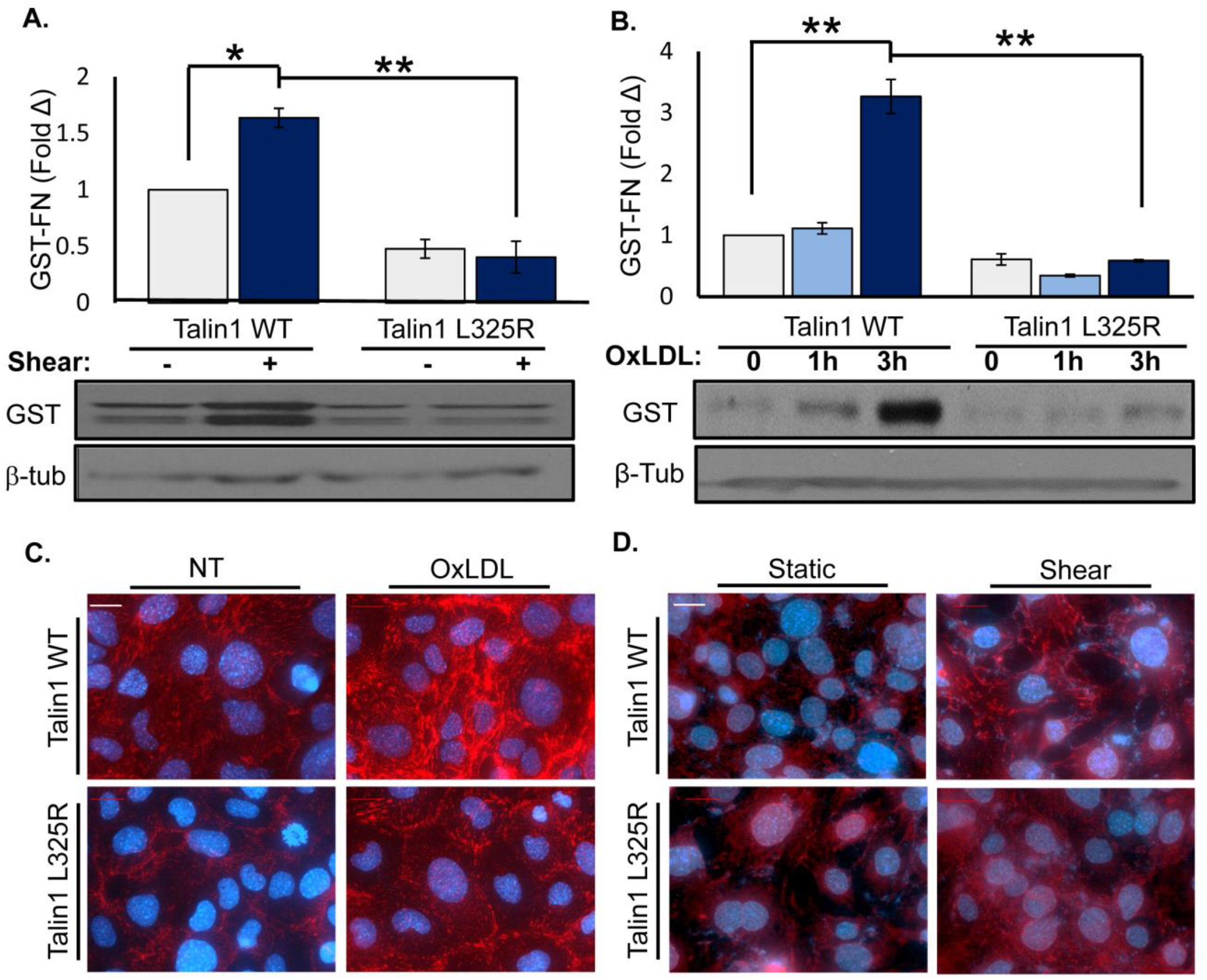
Talin1 Mediates Shear Stress and OxLDL (oxidized low-density lipoprotein)-Induced Integrin Activation. **A/B**) Talin1 WT and Talin1 L325R cells were plated on fibronectin and exposed to flow (5 minutes) or treated with oxLDL (100 ug/ml) for 1 or 3 hours. α5β1 activation was assessed by measuring GST-FNIII_9-11_ retention by western blot. n=4. C/D) Talin1 WT and Talin1 L325R cells were plated on fibronectin, then either treated with oxLDL (C) or exposed to shear stress (D) for 1 hour, then immunostaining for 9EG7 was performed. n=4-5. Values are means ±SE. *P<0.05 and **P<0.01 compared with 0 h time point. 2-way ANOVA with Bonferroni posttest (E-F) were used for statistical analyses.

### Talin1 L325R mutation blocks endothelial inflammation on fibronectin

At atherosclerosis prone regions, NF-κB signaling plays a major role in driving proinflammatory endothelial activation(4). Signaling through α5β1 mediates NF-κB activation and enhanced proinflammatory gene expression in response to both oxLDL(8, 9) and shear stress(26, 27, 32–34). Since integrin inhibitors (small molecule inhibitors, blocking antibodies) blunt shear stress and oxLDL-induced NF-κB activation, we and others have postulated that new integrin-matrix interactions following integrin affinity modulation likely drive endothelial responses to both stimuli. However, these inhibitors do not specifically block high affinity integrins may also compete with stable integrin-matrix interactions. Therefore, we utilized talin1 WT and L325R endothelial cells to study the specific role of integrin affinity modulation in response to shear stress and oxLDL. Whereas both oxLDL and shear stress induce phosphorylation of the NF-κB p65 subunit (hereafter NF-κB), neither oxLDL nor shear stress induced significant NF-κB phosphorylation in talin1 L325R endothelial cells **(Figure 3A/B)**. Similarly, NF-κB nuclear translocation in response to oxLDL and shear stress was observed in talin1 WT but not talin1 L325R endothelial cells (**Figure 3C/D**). In contrast, other known shear-responsive signaling events, such as activation of ERK1/2, AKT, and eNOS (**Supplemental Figure V**) were activated similarly in talin1 WT and talin1 L325R endothelial cells, indicating these signaling responses are independent of shear stress-induced integrin affinity modulation(35). Since α5β1 mediates oxLDL and flow-induced proinflammatory gene expression, we examined the effect of blocking talin1-dependent integrin affinity modulation in this response. Consistent with the suppressed NF-κB activation, our results show a marked inhibition of VCAM-1 and ICAM-1 in response to chronic oscillatory shear stress (OSS) (**Figure 4A/B**) and oxLDL (**Figure 4C/D**) in talin1 L325R endothelial cells. However, talin1-dependent integrin affinity modulation is not required for endothelial VCAM-1 expression in response to TNFα, IL-1β and LPS **(Figure 4E**), consistent with a specific role for integrin signaling in oxLDL and flow-induced proinflammatory endothelial activation.

**Figure 3.**
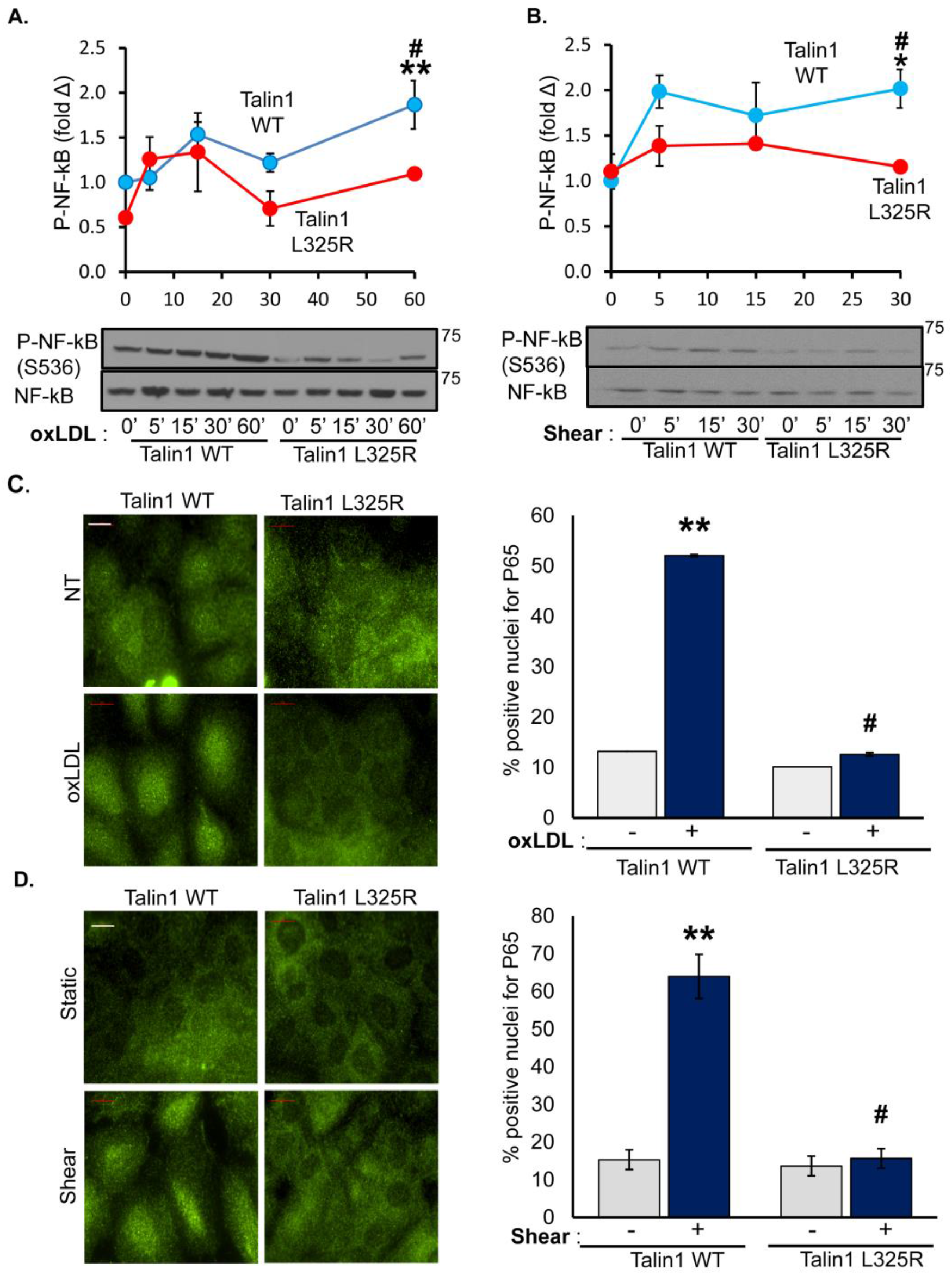
Blocking Talin1-Dependent Integrin Activation Blunts OxLDL (oxidized low-density lipoprotein) and Flow-Induced NF-κB Activation. A/B) Talin1 WT and Talin1 L325R cells were plated on fibronectin and treated with oxLDL or exposed to shear stress for the indicated time points. Western blotting was performed for P-NF-κB (p65, Ser536). Representative western blots are shown (n=4-5). B/D) Talin1 WT and Talin1 L325R cells were treated with oxLDL for an hour or flow for 30 minutes and then fixed and stained for NF-κB and positive nuclear translocation was quantified. Representative images are shown(n=4). D) Talin1 WT and Talin1 L325R cells were exposed to shear stress for the indicated times and P-NF-κB (p65, Ser536) was measured using western blot. Representative western blots are shown (n=4-5). Values are means ±SE. *P<0.05 and **P<0.001 compared *P<0.05 compared with 0 h time point. #P<0.05 compared with Talin1 L325R. 2-way ANOVA with Bonferroni posttest was used for statistical analyses.

**Figure 4.**
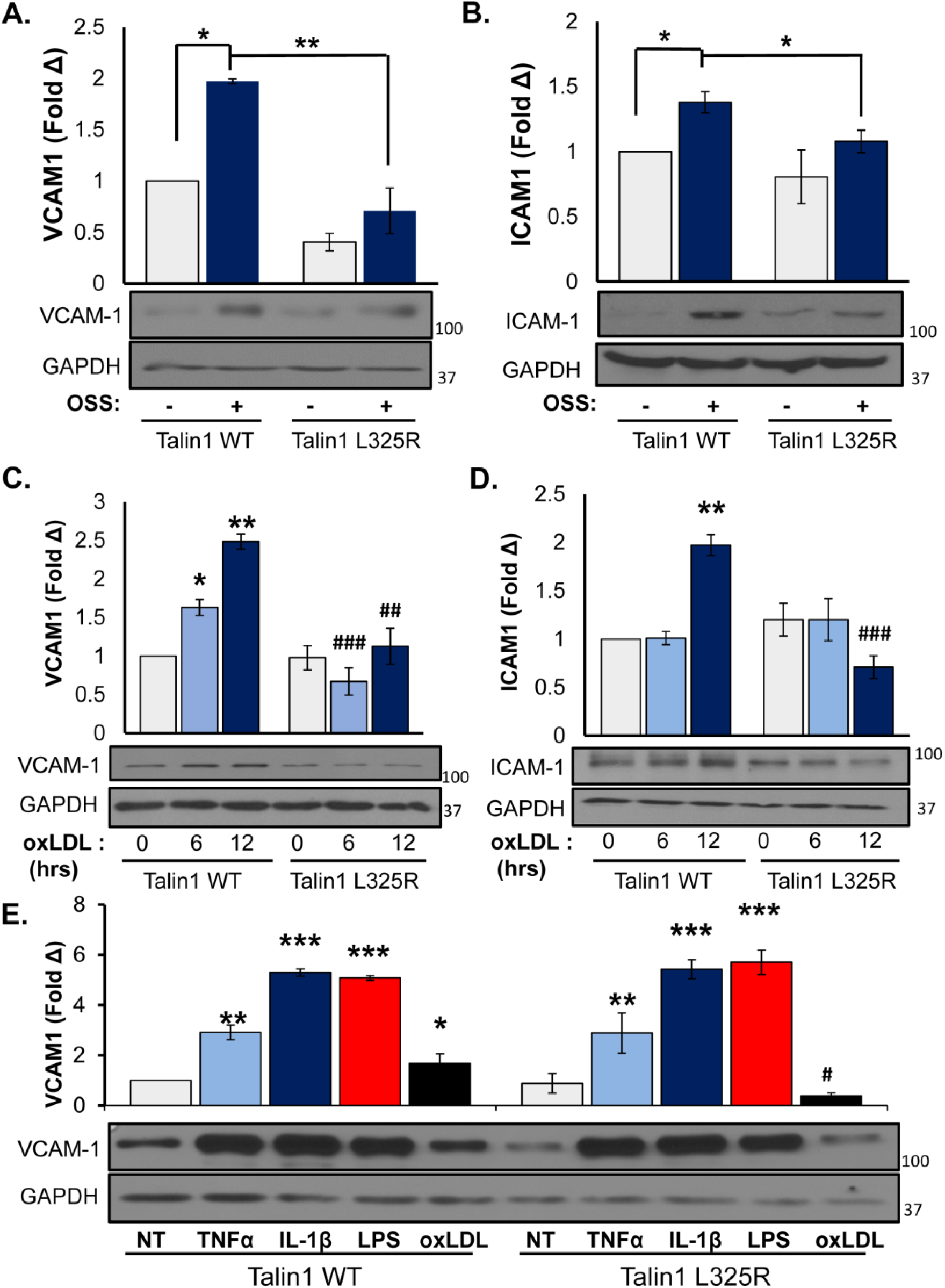
Talin1-Mediated Integrin Activation Is Required for OxLDL (oxidized low-density lipoprotein) and Flow-Induced Inflammation. A-B) Talin1 WT and Talin1 L325R MLECs (mouse Lung endothelial cells) were plated on fibronectin and exposed to oscillatory shear stress (OSS) for 18 hours. VCAM-1(vascular cell adhesion molecule-1) and ICAM-1(intercellular adhesion molecule-1) protein expression was measured using western blot. Representative western blots are shown (n=4-5). C-D) Talin1 WT and Talin1 L325R MLECs were treated with oxLDL for the indicated time points. Immunoblotting was conducted to assess VCAM-1 (C) and ICAM-1 (D) expression. Representative western blots are shown (n=4). D) Talin1 WT and Talin1 L325R MLECs were treated with TNFα (tumor necrosis factor α) (1 ug/ml), IL-1β (interleukin 1β) (5 ug /ml), LPS (lipopolysaccharide) (10 ug /ml), or oxLDL. VCAM-1 expression was assessed using western blot. Representative western blots are shown (n=3). Values are means ±SE. *P<0.05, **P<0.01 ***P<0.001 compared with 0 h time point. ^#^P<0.05, ^##^P<0.01, and ^###^P<0.001 compared with Talin1 L325R. 2-way ANOVA with Bonferroni posttest was used for statistical analyses.

### Aberrant fibronectin fibrillogenesis in endothelial cells expressing Talin1 L325R mutation

Fibronectin deposition occurs early during atherogenesis and contributes to early endothelial activation in part through inducing α5β1 proinflammatory signaling(5, 11, 26, 27, 36). However, α5β1 integrins also regulate fibronectin deposition in response to oxLDL and shear stress(9, 34), and it is unclear whether integrin affinity modulation is required to mediate fibronectin matrix deposition. Therefore, we sought to examine fibronectin deposition in talin1 L325R mutant endothelial cells. Talin1 WT and talin1 L325R endothelial cells were plated on basement membrane proteins for 4 hours and then were exposed to either disturbed flow or oxLDL for 24 hours. Fibronectin deposition into the deoxycholate-insoluble matrix fraction was compared using Western blotting and immunocytochemistry as previously described(9). Talin1 WT endothelial cells showed a low level of fibronectin deposition under unstimulated conditions that increased significantly in response to both oscillatory shear stress (OSS) (**Figure 5A**) and oxLDL (**Figure 5B**). While fibronectin deposition did not differ between talin1 WT and talin1 L325R endothelial cells under unstimulated conditions, talin1 L325R endothelial cells failed to show enhanced fibronectin deposition in response to either stimulus. To gain further insight into this response, the fibronectin matrix was visualized by immunocytochemistry. As previously shown, both OSS and oxLDL increased fibronectin staining in the subendothelial matrix (**Figure 5C/D**), associated with both increased fluorescence intensity for fibronectin (**Figure 5E/F**) and increased fibronectin fibril length (**Figure 5G/H**). While fibronectin staining and fibril length were similar between talin1 WT and talin1 L325R endothelial cells in unstimulated conditions, both OSS and oxLDL failed to enhance fibronectin staining intensity or fibronectin fibril length in talin1 L325R endothelial cells.

**Figure 5.**
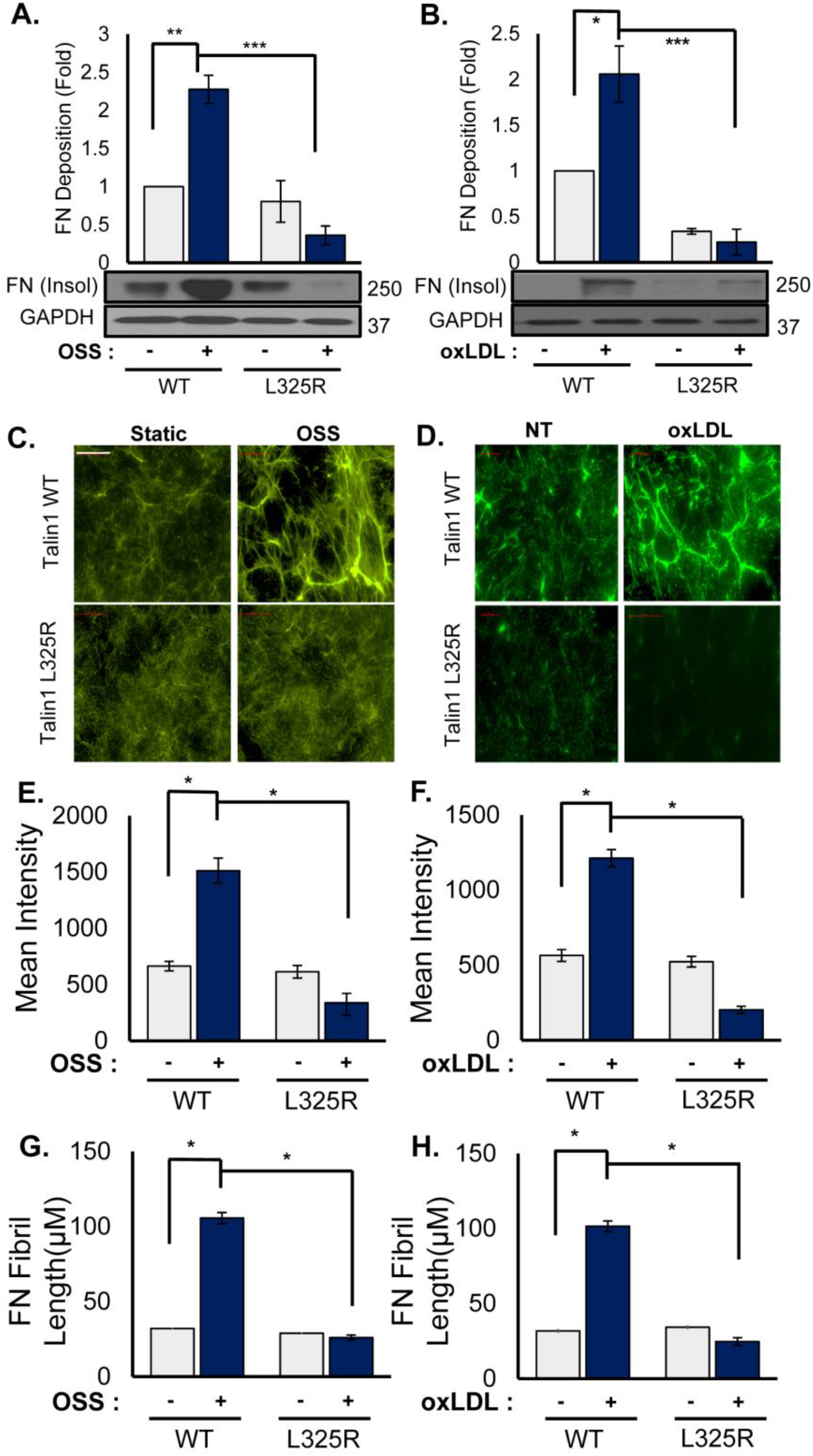
Talin1- Mediated Integrin Activation Promotes Fibronectin Fibrillogenesis. A/B) Talin1 WT and Talin1 L325R MLECs (mouse Lung endothelial cells) were plated on diluted Matrigel for 4 hours and then treated with oxLDL (oxidized low-density lipoprotein) or OSS (oscillatory shear stress) for 24 hours. Deoxycholate insoluble fractions were collected and western blot was conducted for fibronectin.=4. E-H) Talin1 WT and Talin1 L325R MLECs seeded on diluted Matrigel and treated with oxLDL or exposed to OSS for 24 hours. Immunocytochemistry was performed to test fibronectin deposition. Mean fluorescent intensity (E/F) and fibril length (G/H) were assessed. Representative images are shown(n=4-5). Values are means ±SE. *P<0.05, **P<0.01 ***P<0.001 compared with 0 h time point. 2-way ANOVA with Bonferroni posttest was used for statistical analyses.

Although the talin1 L325R endothelial cells were deficient for inducible fibronectin deposition, fibronectin expression was similar between the talin1 WT and talin1 L325R endothelial cells (**Supplemental Figure VI**), suggesting the defect may be in matrix deposition. Fibronectin fibril assembly involves α5β1 translocation from peripheral focal adhesions into tensin1-rich fibrillar adhesions that organize the fibronectin dimers into higher order fibrils(9, 37). To assess whether talin1-dependent integrin affinity modulation affects the formation of fibrillar adhesions in response to oxLDL or OSS, talin1 WT and talin1 L325R endothelial cells were treated with oxLDL or OSS, and the composition of endothelial adhesions was assessed by immunocytochemistry and Western blotting analysis of the isolated focal adhesion fraction. Both oxLDL and OSS stimulated enhanced recruitment of α5 integrins into the focal adhesions in talin1 WT endothelial cells (**Figure 6A/B**), consistent with an increase in fibrillar adhesion formation. However, focal adhesion levels of α5 were unchanged in talin1 L325R endothelial cells. Similarly, both oxLDL and OSS promoted focal adhesion recruitment of tensin1 in talin1 WT but not talin1 L325R endothelial cells (**Figure 6C/D**). Together, these data suggest that inducible high affinity integrins and the formation of new cell-matrix interactions are critical for fibrillar adhesion development and fibronectin deposition in response to oxLDL and OSS in endothelial cells.

**Figure 6.**
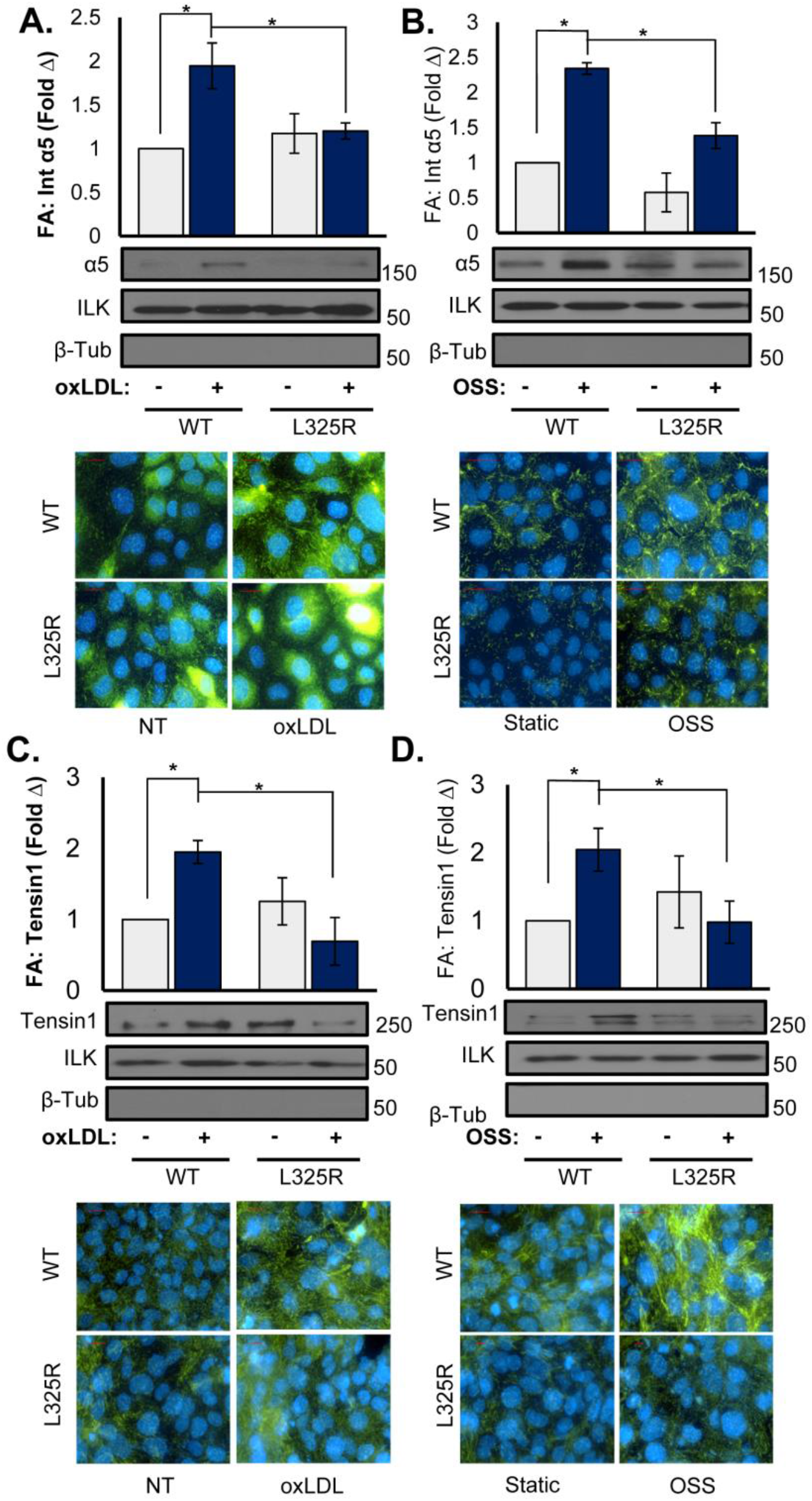
Talin1 L325R Endothelial Cells Show Reduction of α5 and Tenisn1 Localization in Integrin Adhesion in Response to Flow and OxLDL (oxidized low-density lipoprotein). A-B) Talin1 WT and Talin1 L325R cells were treated with oxLDL or exposed to OSS (oscillatory shear stress) for 18 hours and focal adhesions were extracted using hydrodynamic shock or stained for α5 integrins. Representative western blots (=4) and images are shown(n=3-4).C-D) A) Talin1 WT and Talin1 L325R cells were treated with oxLDL or exposed to OSS for 24 hours and focal adhesions were extracted or stained for tenisn1. Representative western blots images are shown(n=4-5). are shown(n=4). Values are means ±SE. *P<0.05 compared with 0 h time point. 2-way ANOVA with Bonferroni posttest was used for statistical analyses.

**Figure 7.**
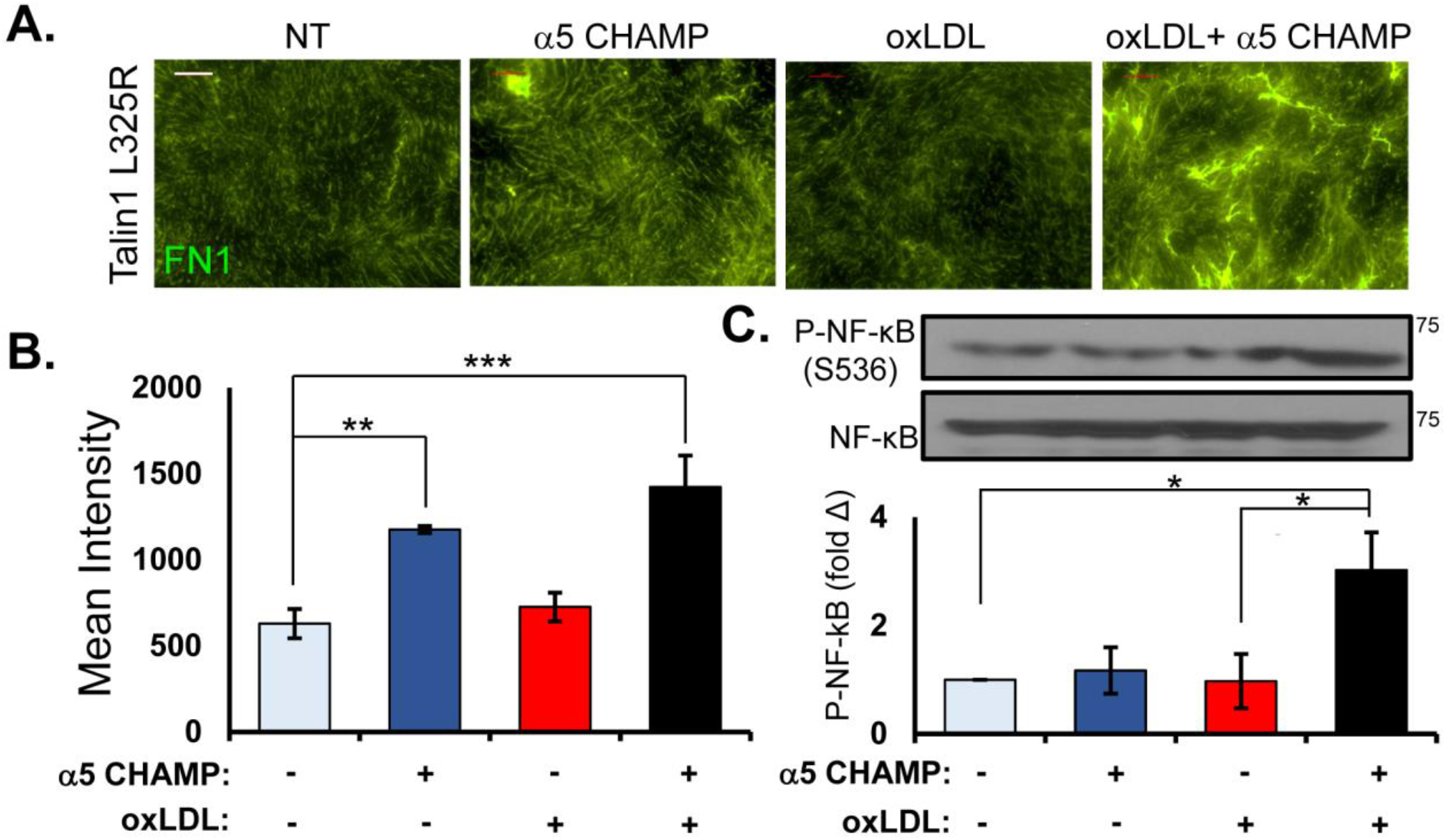
α5 Integrin Activation Requirement is Different for Matrix Remodeling Versus Inflammation. A)Talin1 L325R cells were plated on Matrigel for 4 hours, then treated with either oxLDL, α5 CHAMP, or both for 24 hours. Deoxycholate insoluble fractions were collected and immunocytochemistry was performed for fibronectin and quantified for mean fluorescent intensity. Representative images are shown(n=4-7). C. Talin1 L325R cells were plated on fibronectin and then treated with either oxLDL, α5 CHAMP, or both for 1 hours. Western blot was done for P-NF-κB (p65, Ser536). Representative blots are shown (n=4). Values are means ±SE. *P<0.05, **P<0.01 ***P<0.001 compared with 0 h time point. #p<0.05 compared with oxLDL group. two-way ANOVA with Bonferroni posttest was used for statistical analyses.

### Requirement of integrin signaling for endothelial activation and matrix remodeling

While these data indicate that a subset of endothelial responses to oxLDL require inducible high affinity integrins, it remains unclear whether integrin affinity modulation and signaling are sufficient for these responses. To test the role of integrin affinity modulation in endothelial proinflammatory gene expression and fibronectin deposition, we employed CHAMP peptides targeted to the α5 integrin transmembrane region to selectively induce high affinity α5β1 integrins in the talin1 L325R cells. Treatment with the α5 CHAMP peptide was partially sufficient to induce fibronectin deposition in talin1 L325R endothelial cells **(Figure 6A/B)**, which was fully recovered by treatment with a combination of α5 CHAMP peptides and oxLDL **(Figure 6A/B)**. While α5 is clearly required for oxLDL-induced NF-κB activation(8), activation of α5β1 with the α5 CHAMP peptide was not sufficient to induce NF-κB activation **(Figure 6C)**. However, simultaneous treatment with both oxLDL and the α5 CHAMP peptide restored NF-κB activation in talin1 L325R endothelial cells **(Figure 6C)**, suggesting that oxLDL-induced NF-κB activation requires both integrin-dependent and integrin-independent signaling responses. Taken together, these data suggest that oxLDL-induced α5β1 integrin activation alone can mediate fibronectin assembly, whereas NF-κB activation and maximal fibronectin assembly requires co-stimulatory integrin-dependent and integrin-independent signaling pathways.

### Mice expressing endothelial talin1 L325R are protected from atherogenic inflammation

Since our *in vitro* studies show a clear role for talin1-dependent integrin affinity modulation in atherogenic endothelial activation, we next examined whether mice expressing only talin1 WT or talin1 L325R in their endothelial cells show differential susceptibility to disturbed flow-induced endothelial activation *in vivo*. Tamoxifen-inducible, endothelial-specific talin1 WT (iEC-Talin1 WT; Talin1^WT/flox^, VE-Cadherin-CreERT^tg/?^) and talin1 L325R (iEC-Talin1 L325R; Talin1^flox/L325R^, VE-Cadherin-CreERT^tg/?^) mice underwent partial ligation of the left carotid artery to induce low, oscillatory flow. The right carotid artery remained exposed to laminar flow and was used as an internal control. After 48 hours, intimal mRNA was isolated to analyze endothelial proinflammatory gene expression using qRT-PCR. Intimal mRNA preparations were shown to be enriched for CD31 and deficient in smooth muscle actin compared to medial mRNA (**Figure 8A**). Both iEC-Talin1 WT and iEC-Talin1 L325R mice show reduced KLF2 expression in the left carotid artery following partial ligation, consistent with a loss of laminar flow (**Figure 8B**). However, only iEC-Talin1 WT mice showed a significant increase in proinflammatory gene expression (ICAM-1, VCAM-1) in the left carotid, whereas iEC-Talin1 L325R mice showed no upregulation of proinflammatory gene expression (**Figure 8B**). In addition to mRNA analysis, we harvested carotid arteries from iEC-Talin1 WT and iEC-Talin1 L325R mice 7 days after partial carotid ligation for immunohistochemical analysis. Consistent with our *in vitro* data, iEC-Talin1 L325R mice show a remarkable reduction of endothelial NF-κB nuclear translocation (**Figure 8C/D**), VCAM-1 expression (**Figure 8E/F**), and macrophage (Mac2-positive) recruitment (**Figure 8G/H**). In contrast, carotid endothelial (CD31-positive) and smooth muscle (smooth muscle actin-positive) content was similar between iEC-Talin1 WT and iEC-Talin1 L325R mice (**Supplemental Figure VII**). Altogether, these data show that modulating integrin affinity in endothelial cells contributes to proinflammatory endothelial activation both *in vitro* and *in vivo*.

**Figure 8.**
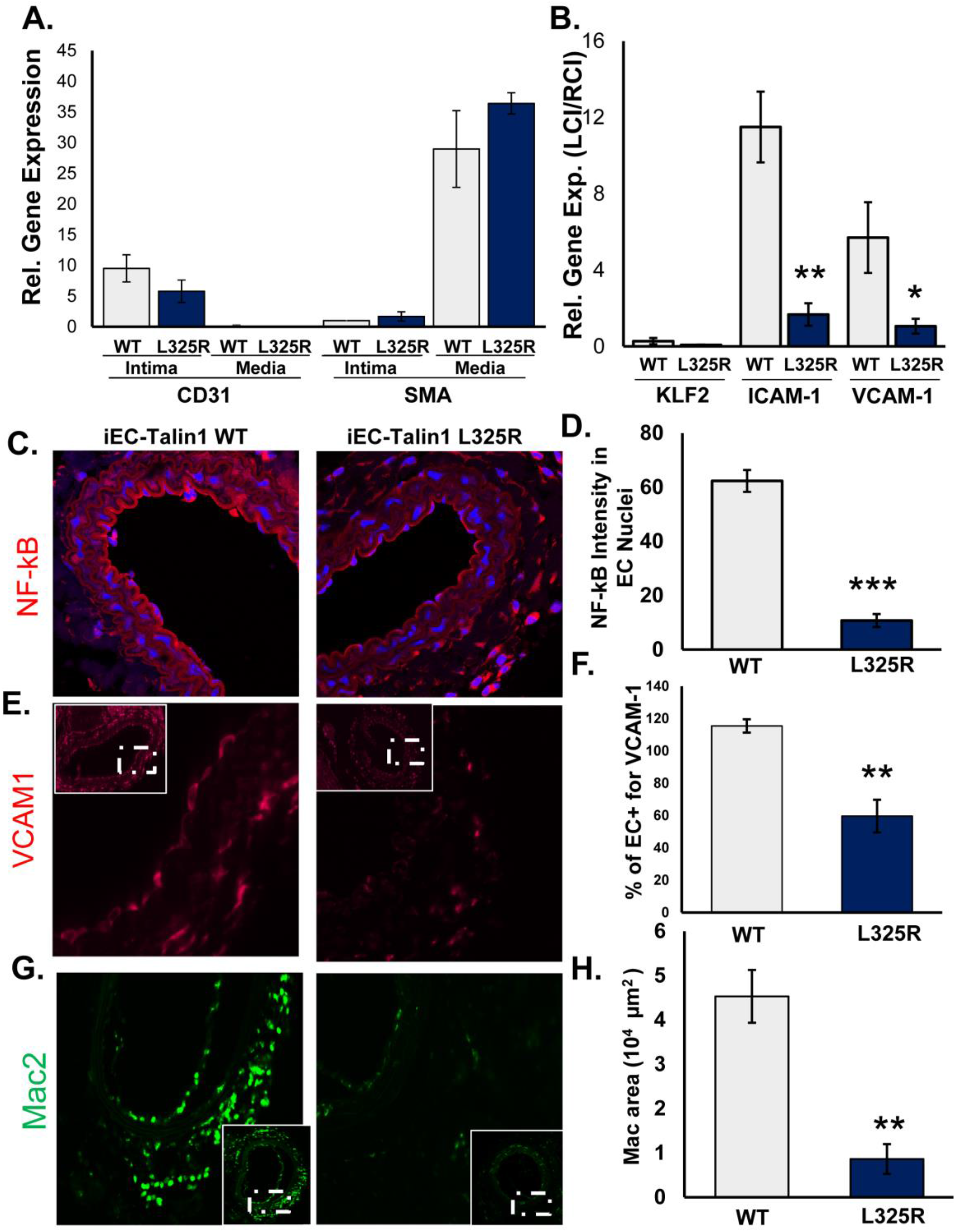
Mice Expressing Endothelial Talin1 325R Mutation Are Protected from Atherogenic Inflammation. **A/B)** iEC (inducible, endothelial cell-specific)-Talin1 WT (Talin1+/flox VE [vascular endothelial]-cadherin-CreERT2tg/?) and iEC Talin1 L325R -(Talin1flox/L325R VE-Cadherin- CreERT2tg) mice were treated with tamoxifen for 5 consecutive days. Four weeks later, mice underwent partial carotid ligation (PCL) of their left carotid artery for 48 hrs and the endothelial mRAN was isolated. Expression of CD31, SMA (A), Klf2, ICAM-1, and VCAM-1 (B) was determined by quantitative real-time PCR. n=5-6. **C-H)** iEC-Talin1 WT and iEC Talin1 L325R were undergone PCL for 7 day. Tissues were collected and immunohistochemistry for NF-κB (visualized by confocal microscopy), VCAM1, or Mac2 was conducted. N=5-6 for each group. F-H) After PCL for 7 days, immunohistochemistry was performed for MAC2(green) and SMA (red). Representative images are shown(n=5-6). Values are means ±SE. *P<0.05 and ***P<0.001 compared with 0 h time point. n.s. indicates not significant. Student t test was used for statistical analyses.

## Discussion

Cell-matrix interactions play a critical role in endothelial activation and early atherosclerosis(2), however, the role of integrin affinity modulation in this context remains enigmatic. Previously, we showed that fibronectin enhances proinflammatory responses to flow and oxLDL and deleting fibronectin-binding integrins in the endothelium blunts atherogenic inflammation(8, 9). Using mice expressing the L325R Talin1 mutation that selectively inhibits integrin affinity modulation without affecting talin1’s ability to link the integrin β tail with actin, we demonstrate that talin1-dependent integrin affinity modulation is crucial for endothelial proinflammatory activation in response to atherogenic stimuli. Consistent with cell lacking talin1(31), talin1 L325R endothelial cells adhere and form focal adhesions but fail to form nascent adhesions at the edge of membrane protrusion resulting in a reduction in endothelial cell spreading. However, talin1 L325R endothelial cells form stable adhesions that do not differ significantly in size or number from talin1 WT endothelial cells under unstimulated conditions. Additionally, we showed that talin1-dependent α5β1 affinity modulation in response to oxLDL and flow is critical for fibronectin deposition, NF-κB activation, and proinflammatory gene expression. Lastly, mice harboring a talin1 L325R mutation in endothelial cells show a significant reduction in proinflammatory gene expression and macrophage accumulation in the partial carotid ligation model of disturbed flow, suggesting that induction of high affinity α5β1 integrins in endothelial cells critically regulates atherogenic endothelial activation.

While talin1 critically regulates cell adhesion in leukocytes and platelets, multiple mechanisms may mediate cell adhesion and spreading in talin1 L325R endothelial cells. The presence of talin1 L325R in endothelial cells may allow the endothelial cell integrins to attain the extended, closed (intermediate affinity) conformation, which could mediate weak integrin-matrix interactions to support initial cell adhesion and spreading in the mild mechanical environment of static cell culture. However, talin1 L325R cells show significantly reduced nascent adhesion formation, suggesting that these intermediate affinity integrins may not be sufficient to stabilize membrane protrusions to support nascent adhesion formation. Consistent with this hypothesis, low levels of force application to extended, closed integrins may promote integrins to attain the extended, open conformation and stabilize this high affinity conformation through a process termed outside-in integrin activation(38). This method of integrin activation may be insensitive to talin1-dependent transition to a high affinity state in the context of mechanical coupling. Nascent adhesions form in a tension-independent manner and may be insensitive to outside-in affinity modulation through mechanical coupling(39), whereas outside-in integrin activation may support focal adhesion remodeling to force in the absence of inducible affinity modulation.

Like our results in endothelial cells, talin1-deficient fibroblasts adhere and spread normally(31), and siRNA-mediated knockdown of talin2 in these cells does not blunt initial cell adhesion and spreading. However, in the absence of talin2, the cells fail to form stable adhesions and ultimately lose the spread cell phenotype(31). Expression of the talin1 head domain in these cells restores nascent adhesion formation, but only rescue with full length talin1 rescued focal adhesion formation, suggesting that tension-independent integrin activation mediates nascent adhesion formation whereas mechanical coupling through talin1-actin interactions critically regulates focal adhesion formation. Compared to fibroblasts, endothelial cells express minimal talin2(22, 40), and deletion of endothelial talin1, but not talin2, results in defective cardiovascular development(29, 41). While talin1 knockdown reduces cell adhesion and focal adhesion formation, rescue with either talin1 or talin2 is sufficient to rescue both adhesion and focal adhesion formation, whereas expression of W359A and L325R talin did not(22). In contrast, our data suggest that talin1 L325R mutant endothelial cells show normal focal adhesion formation but impaired α5β1 affinity modulation in response to oxLDL or flow, indicating that endothelial talin2 expression is not sufficient to support inducible integrin affinity modulation in endothelial cells. Furthermore, talin1, but not talin2, is preferentially recruited to protrusions and localizes to peripheral adhesions, suggesting that talin2 likely plays a minimal role in nascent adhesion formation during cell motility and spreading(42).

Integrins are involved in a vast spectrum of biological processes, including cell adhesion, gene expression, proliferation, and cell migration(43). Deletion of fibronectin or fibronectin-binding integrins (α5β1, αvβ3) limits endothelial NF-κB activation and proinflammatory gene expression in atherosclerosis models(9, 11), whereas β1 integrin inhibition reduces vascular permeability in response to proinflammatory mediators (LPS, IL-1β)(44). To dissect the role of integrin signaling in endothelial cell phenotype, several groups have used variety of integrin inhibitors and blocking antibodies to implicate integrin affinity modulation and new integrin-matrix interactions in endothelial proinflammatory responses, particularly in response to flow and oxLDL(8, 12). However, integrin inhibitors and blocking antibodies have the potential to affect pre-existing integrin-matrix interactions as well, making it difficult to determine the specific role of integrin affinity modulation and new integrin-matrix interactions. In addition, the use of ligand-mimetic inhibitors may stimulate ligation-dependent signaling, and anti-integrin antibodies may induce a subset of integrin signaling due to clustering. Our model of talin1 L325R mutant endothelial cells eliminates these barriers by specifically eliminating talin1-dependent integrin affinity modulation without affecting talin1-dependent focal adhesion formation. The talin1 L325R mutant endothelial cells provide the first conformation of an important role for endothelial integrin affinity modulation in fibronectin deposition, NF-κB activation, and proinflammatory adhesion molecule expression (VCAM-1, ICAM-1). Furthermore, we utilize α5β1 activating CHAMP peptides to show that α5β1 affinity modulation is partially sufficient to induce fibronectin deposition, similar to previous studies by Wu et al. using constitutively active integrins and integrin activating antibodies(45). However, CHAMP peptide-induced high affinity α5β1 integrins were not sufficient to induce maximal fibronectin deposition and NF-κB activation, suggesting that both integrin-dependent and integrin-independent signaling pathways contribute to these responses.

Previous studies in mouse models clearly implicate cell-matrix interactions in atherogenic endothelial activation but fail to connect endothelial integrin affinity modulation to these responses. The endothelial-specific talin1 L325R mutant mouse is the first model that directly tests the role of endothelial integrin affinity modulation *in vivo*. Interestingly, endothelial talin1 knockout mice show destabilized endothelial cell-cell junctions, mostly due to disruption of vascular endothelial-cadherin organization. While talin1 reexpression restores endothelial adherens junction structure, rescuing integrin activation with the talin1 head domain (but not a L325R head domain mutant) only partially reverts this phenotype. These data suggest an important role for both talin1-dependent integrin affinity modulation and talin1-mediated mechanical coupling in maintaining vascular barrier function(30). Similarly, β1 integrin expression is crucial for the development of stable, non-leaky vessels(46). However, inhibiting β1 integrins limits LPS-induced vascular leakage, suggesting that integrin signaling may play differential roles in regulating vascular permeability in developing blood vessels compared to mature vessels.

Much of our current understanding of integrin affinity modulation derives from studies in platelets and leukocytes, leading to the development of a multiple therapeutics for hematological and cardiovascular diseases. However, far less is known concerning integrin affinity modulation in adherent cell types, such as the endothelium. These studies demonstrate for the first time a vital role for endothelial integrin affinity modulation in the regulation of endothelial phenotype and proinflammatory gene expression *in vitro* and *in vivo*. Specifically, we provide the first evidence that talin1-dependent induction of high affinity α5β1 integrins drives endothelial proinflammatory responses *in vitro*, and selectively blocking the induction of high affinity integrins in endothelial cells reduces endothelial proinflammatory responses to disturbed flow *in vivo*. Through a better understanding of the molecular mechanisms regulating endothelial integrin affinity modulation, we may be able to generate future therapeutics that limit endothelial activation to reduce inflammation in a variety of pathological contexts.

## Materials and Methods

### Cell culture

Mouse lung endothelial cells were isolated from Talin1^fl/L325R^ mice (gift of Brian Petrich, Emory, Atlanta, GA) as previously described(30). Briefly, lung tissue was harvested, minced, and pushed through a 16G needle and subjected to enzymatic digestion to obtain a single cell suspension. After sorting with magnetic beads coupled to ICAM2 antibodies (eBiosource), cells were transformed using a retrovirus expressing temperature-sensitive large T-antigen. The floxed allele of talin1 was deleted using adenovirus harboring either GFP (green fluorescence protein)-Cre or GFP alone and then sorted for GFP positivity. Lung endothelial cells were grown in DMEM with 10% fetal bovine serum, 1% penicillin/streptomycin, and 1% glutamax. For endothelial shear stress treatments, cells were plated on 38 × 75 mm glass slides (Corning) at confluency, and slides were assembled into a parallel plate flow chamber as previously described(47). For acute shear stress, cells were exposed to 12 dynes/cm2 flow for up to 1 hour. For chronic oscillatory flow (model of disturbed flow), cells were exposed to ±5 dynes/cm2 with 1 dyne/cm2 forward flow for media exchange. Human LDL (Intracell) was oxidized by dialysis in PBS containing 13.8 μM Cu2SO4 followed by 50 μM EDTA as previously described(28). Computationally designed transmembrane α-helical peptides (CHAMP) peptides were a gift of Dr. William DeGrado (University of California – San Francisco)(48).

### Western blot

Cell lysis and western blot was done as previously described(28). Briefly, cells were lysed in 2X laemmli buffer, separated by SDS-PAGE gels and then transferred to polyvinylidene difluoride (PVDF) membranes (Bio-Rad, Hercules, CA). Consequently, membranes were blocked in 5% dry milk in TBST for an hour, then incubated with primary antibodies overnight. Antibodies used include rabbit anti-P-p65 (Ser 536), rabbit anti-β-Tubulin, rabbit anti-GAPDH, rabbit anti-FAK (Tyr 397) (Cell Signaling Technologies), rabbit anti-integrin α5, rabbit anti-integrin αv, rabbit anti-ILK, rabbit anti-VCAM1, rabbit anti-ICAM1, rabbit anti-β1 integrin (9G7) (BD Pharmingen) (**Supplemental Table I**). The following day after rinsing with TBST, HRP-conjugated secondary antibodies (Jackson ImmunoResearch) in blocking buffer were applied for two hours. Antibodies were detected using Pierce ECL solution (ThermoFisher) and x-ray film (Phenix Research Products).

### Immunocytochemistry

Cells were plated on coverslips or slides and fixed in formaldehyde and permeabilized using 0.2% Triton X100 solution. Coverslips-containing cells were blocked with 10% horse serum for an hour. Subsequently, cells were washed and incubated with primary antibodies overnight (**Supplemental Table I**). Next, fluorochrome-tagged secondary antibodies (Life Technologies) were added to coverslips for 2 hours. After rinsing, cells were counterstained with DAPI. We used Nikon Eclipse Ti inverted epifluorescence microscope equipped with a Photometrics CoolSNAP120 ES2 camera and the NIS Elements 3.00, SP5 imaging software.

### Adhesion/spreading assay

Cells were plated sparsely on fibronectin-coated coverslips for 15, 30, or 60 minutes in serum-free DMEM. Next, cells were fixed with PBS-buffered, 4% formaldehyde and co-stained with 546-conjugated phalloidin and 9EG7 (active β1 integrin).

### Integrin Activation Assay

Activation of α5β1 integrins was assessed using a specific α5β1 ligand mimetic as previously described(8). Briefly, after cells were stimulated, a glutathione S-transferase (GST) tagged portion of the FN protein (FN III 9-11) was added to the cell culture medium for 30 minutes. Cell were rinsed with fresh media and then lysed with 2X laemmli buffer, subjected to western blotting, and probed with rabbit anti-GST (~70kDa).

### Focal Adhesion Isolation

Endothelial cells were plated on glass slides and treated. After experiments, cells were exposed to hypertonic shock using triethanolamine (2.5 mmol/L at pH 7.0) for 3 minutes. Next, cell bodies were removed by pulsed hydrodynamic force (Conair WaterPIK) at ≈0.5 cms from and ≈90° to the surface of the slide scanning the entire length 3 times. Slides were visualized under a microscope to ensure complete removal of cell bodies before lysis in 2X Laemmli buffer.

### Deoxycholate Solubility Assay

Fibronectin integrated into extracellular matrix is insoluble in a deoxycholate detergent. Discerning between the two pools of fibronectin (intracellular vs matricellular, ie. extracellular) was done using a deoxycholate solubility assay, as previously described(9). Briefly, after stimulation, cells were rinsed with ice cold PBS, incubated with DOC buffer (2% Sodium Deoxycholate, 20mM Tris-HCL, pH 8.8, 2 mM PMSF, 2mM Iodoacetic Acid (IAA), 2mM N-ethylmaleimide (NEM), 10mM EDTA, pH 8), lysed, and then passed through a 25G needle. The DOC-insoluble matrix fraction was pelleted by centrifugation, and the DOC-soluble supernatant fraction was collected. After rinsing the pellet with additional DOC buffer, the DOC-insoluble faction was lysed in a solubilization buffer (2% SDS, 25mM Tris-HCL, pH 8, 2mM PMSF, 2mM Iodoacetic Acid (IAA), 2mM N-ethylmaleimide (NEM), 10mM EDTA, pH 8). Samples were subjected to western blotting and probed with rabbit anti-fibronectin.

### Quantitative real-time PCR

qRT-PCR was done as previously described(12). Briefly, mRNA was extracted from tissues using TRIzol (Life Technologies, Inc., Carlsbad, CA). Next, iScript cDNA synthesis kit (Bio-Rad, Hercules, CA) was used to synthesize Complimentary DNA. qRT-PCR was done using Bio-Rad iCycler with the use of SYBR Green Master mix (Bio-Rad). We used the online Primer3 software and verified by sequencing the PCR products (**Supplemental Table I**). Results were expressed as fold change by using the 2-ΔΔCT method.

### LDL Oxidation

Low density lipoprotein was purchased from Alfa Aesar and oxidized using 13.8 mM copper sulfate as previously described(8). Briefly, LDL was dialyzed in 1xPBS for 24 hours, then oxidized in 1xPBS containing 13.8 <u>mM</u> copper sulfate for 24 hours followed with 50 μmol/L EDTA in 1x PBS for another day. Nitrogen gas was infused over oxLDL prior to storage, and oxLDL was tested for endotoxin contamination using a chromogenic endotoxin quantification kit (Thermo Scientific).

### Animals and Tissue Harvesting

The Louisiana State University Health Sciences Center-Shreveport Animal Care and Use Committee approved all animal protocols, and all animals were cared for according to the National Institutes of Health Guide for the Care and Use of Laboratory Animals. C57Bl/6J mice with a tamoxifen inducible, VE-cadherin-CreERT2 transgene (from Ralf Adams, Max Planck, Germany) with either wildtype talin1, floxed alleles for talin1, or with an L325R mutation in talin1(49). All mice were backcrossed to C57BL/6J mice for at least 7 generations. At 8 weeks of age, mice were treated with 1 mg/kg tamoxifen (Sigma-Aldrich, St Louis, MO) via intraperitoneal injection every day for 5 total injections to induce Cre expression and gene excision. Four weeks after tamoxifen injection, mice underwent partial carotid ligation as previously described(50). Briefly, a superficial midline incision on the neck of mice under isoflurane anesthesia was done to expose the left carotid artery. Two 7-0 silk sutures were used to tie-off (occlude flow) to the internal, external, and occipital branches of the left carotid artery, whereas flow remained patent through the superior thyroid artery branch. The incision was closed with surgical glue and 6-0 silk sutures. Carprofen (5mg/kg) was administered as a post-surgical analgesic and 800uL of saline solution was given subdurally to prevent dehydration stress. Surgical success was confirmed via echocardiography 1 to 2 days prior to endpoint for confirmation of ligation. Mice were euthanized by pneumothorax under isoflurane anesthesia for tissue collection either 2- or 7-days post-surgery. After 2 days, carotid arteries were collected for RNA isolation performed by a TRIzol flush as previously described(50). Briefly, carotids were cleaned of perivascular adipose tissue and flushed with 150 mL TRIzol from an insulin syringe. The remaining media/adventitia were then placed in 150 mL TRIzol and sonicated to lyse the tissue. Samples were then frozen until analysis by quantitative PCR. After 7 days, the carotids were excised, placed in 4% PBS buffered formaldehyde, and processed for immunohistochemistry.

### Immunohistochemistry

Tissue was fixed in PBS-buffered, 4% formaldehyde, processed for paraffin embedding, and cut into 5 μm thick sections onto superfrost plus glass sides. After heated and subjected sodium citrate antigen retrieval (Vector Labs, H-3300), tissue was blocked using 10% horse serum, 1% BSA in PBS. Primary antibodies were applied overnight, followed by Alexa Fluor conjugated secondary antibodies (Thermo Fisher). Stains are imaged on a Nikon Eclipse Ti inverted fluorescent microscope. Images are captured at either a 10X, 20X, or 60X (oil objective) using the Photometrics Coolsnap120 ES2 camera and the NIS Elements BR 3.00, SP5 imaging software. NF-κB nuclear translocation was visualized by Leica TCS SP5 confocal microscope using 63X (oil objective) and the Nikon NIS-Elements C software.

### Statistical analysis

Statistical analysis was performed using GraphPad Prism software. All of the data were tested for normality using Kolmogorov-Smirnov test. Data passing the normality tests were analyzed using either Student’s t-test, 1-way ANOVA with Newman-Keuls post-test, or 2-way ANOVA with Bonferroni post-tests depending upon the number of independent variables and groups. Data failing the normality test were analyzed using the nonparametric Mann-Whitney U test and the Kruskal-Wallis test with post hoc analysis.

## Acknowledgements

None.

---

**Supplemental Figure I.**
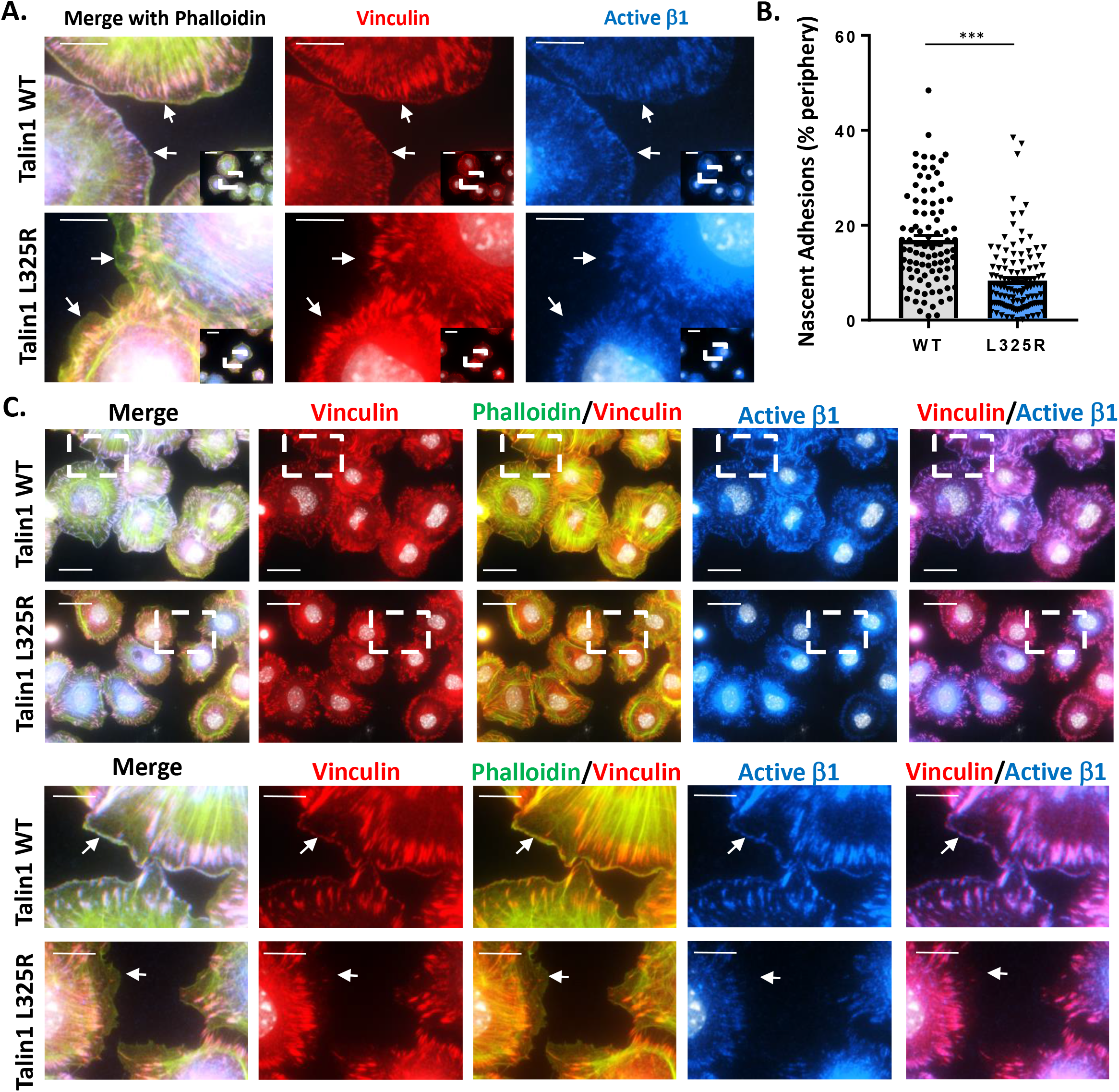
Talin1 L325R Mutation Limits Nascent Adhesion Formation. Talin1 WT and Talin1 L325R cells were plated on fibronectin for 30 minutes and stained for actin (phalloidin, green), vinculin (red), and active (high affinity) β1 (blue). A. Representative micrographs are shown with 60x inset. Scale bars = 10 μm and 20 μm in inset. B. Nascent adhesions were identified as vinculin and β1 integrin positive within 1 μm of the cell edge. Percent of the cell periphery positive for nascent adhesions (active β1) were quantified for individual cells from each of three independent experiments. C. Addition images of altered nascent adhesion formation between talin1 WT and talin1 L325R endothelial cells. Representative micrographs are shown with 60x inset. Scale bars = 10 μm and 20 μm in inset. Representative micrographs from one of three independent experiments are shown.

**Supplemental Figure II.**
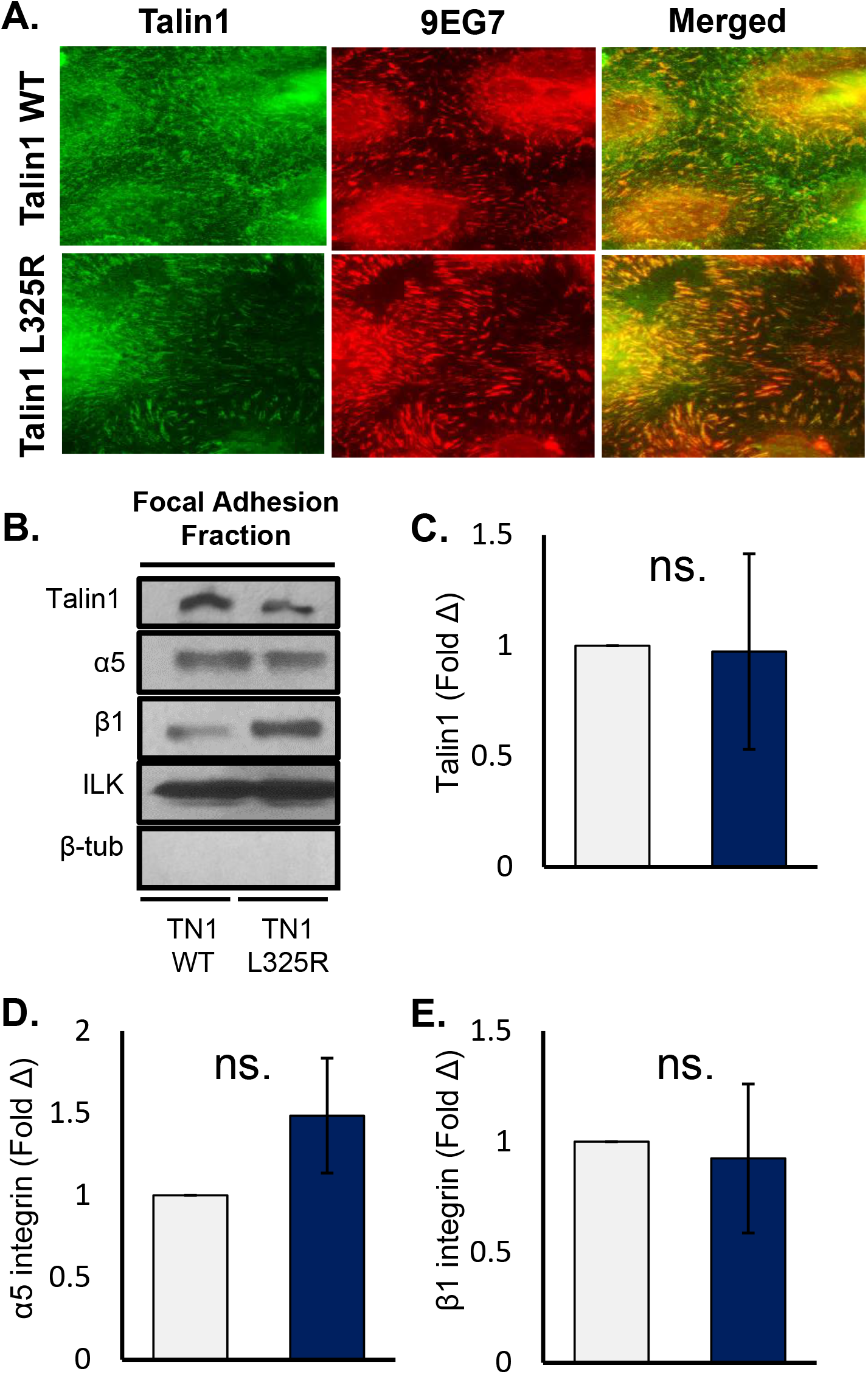
Talin1 L325R Mutation Does Not Alter Focal Adhesion Structure. A) Talin1 WT and Talin1 L325R MLECs were plated on fibronectin, then fixed and immunostained for Talin1 and 9EG7 (active β1). Representative images are shown(n=3). B) Talin1 WT and Talin1 L325R cells were plated on fibronectin, then focal adhesions were extracted. Representative western blots are shown, and quantification of talin1, α5, and β1 levels are indicated. (n=3).

**Supplemental Figure III.**
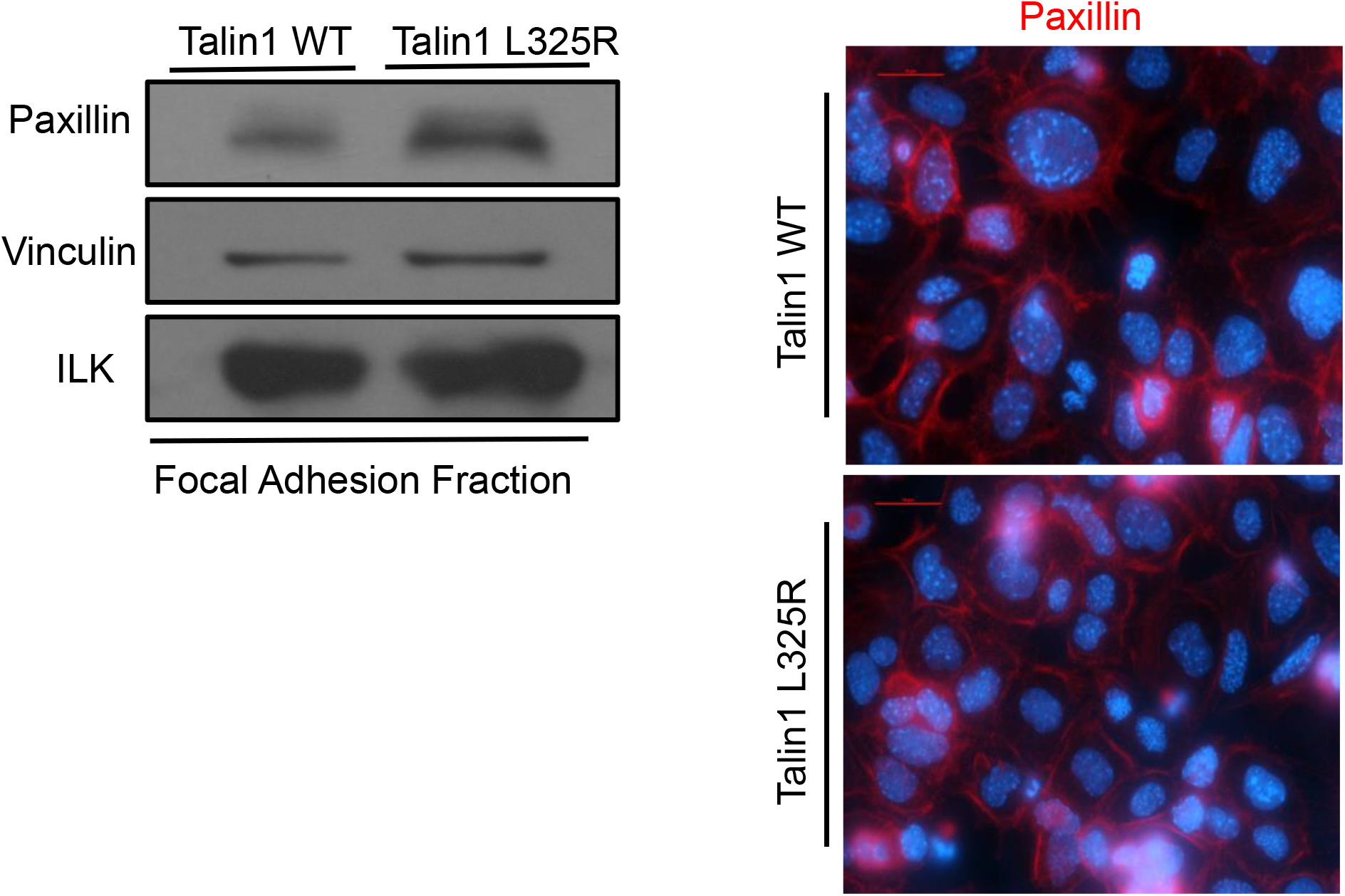
Talin1 L325R Mutation Does Not Alter Focal Adhesion Structure. Talin1 WT and Talin1 L325R cells were plated on fibronectin, then focal adhesions were extracted, or cells fixed for immunocytochemistry. Representative western blots and images are shown (n=3).

**Supplemental Figure IV.**
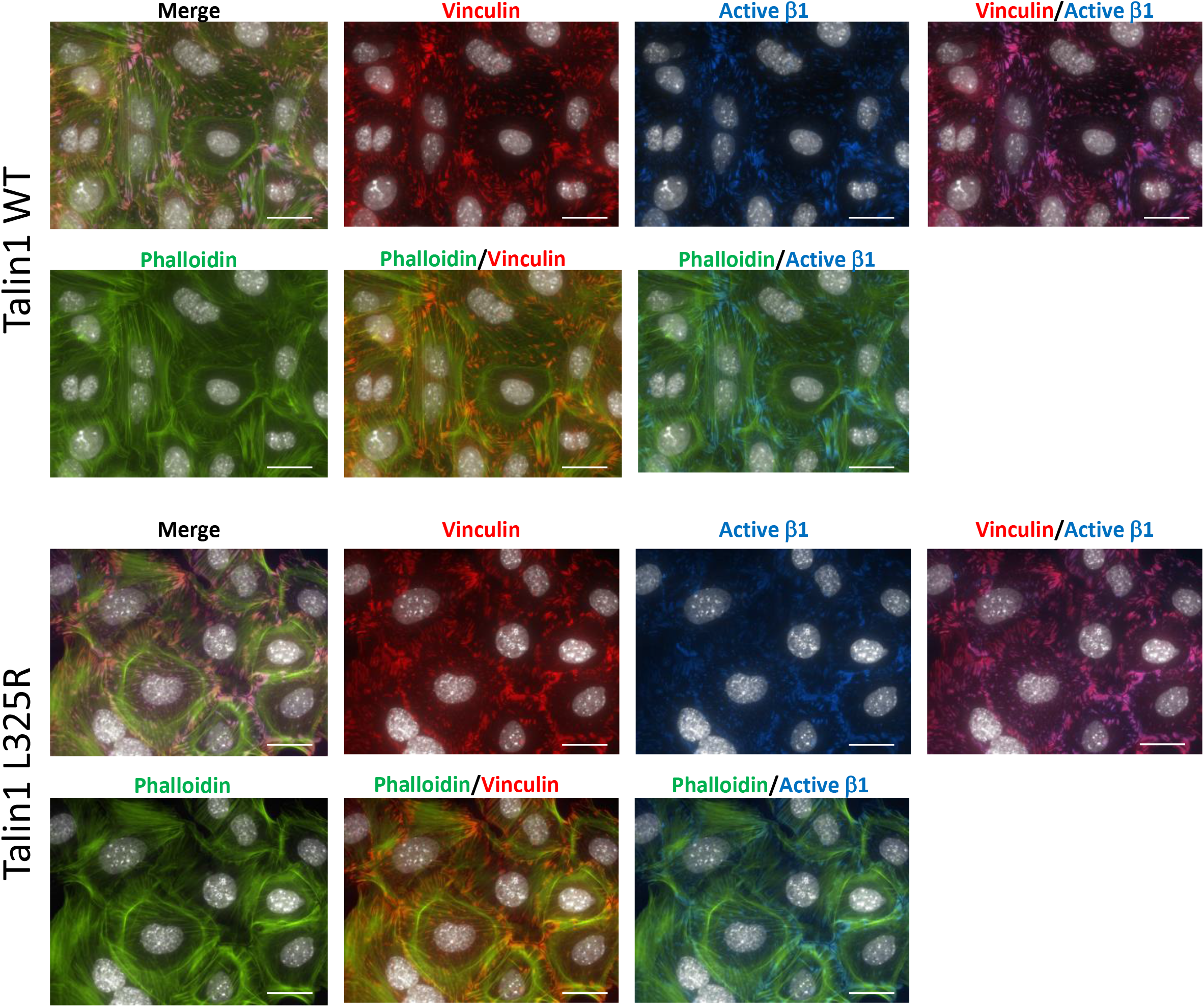
Talin1 WT and Talin1 L325R Endothelial Cells Show Similar Focal Adhesion Formation. Talin1 WT and Talin1 L325R cells were plated on fibronectin for 120 minutes and stained for actin (phalloidin, green), vinculin (red), and active (high affinity) β1 (blue). Representative 60X micrographs are shown. Scale bars = 20 μm. Representative micrographs from one of three independent experiments are shown.

**Supplemental Figure V.**
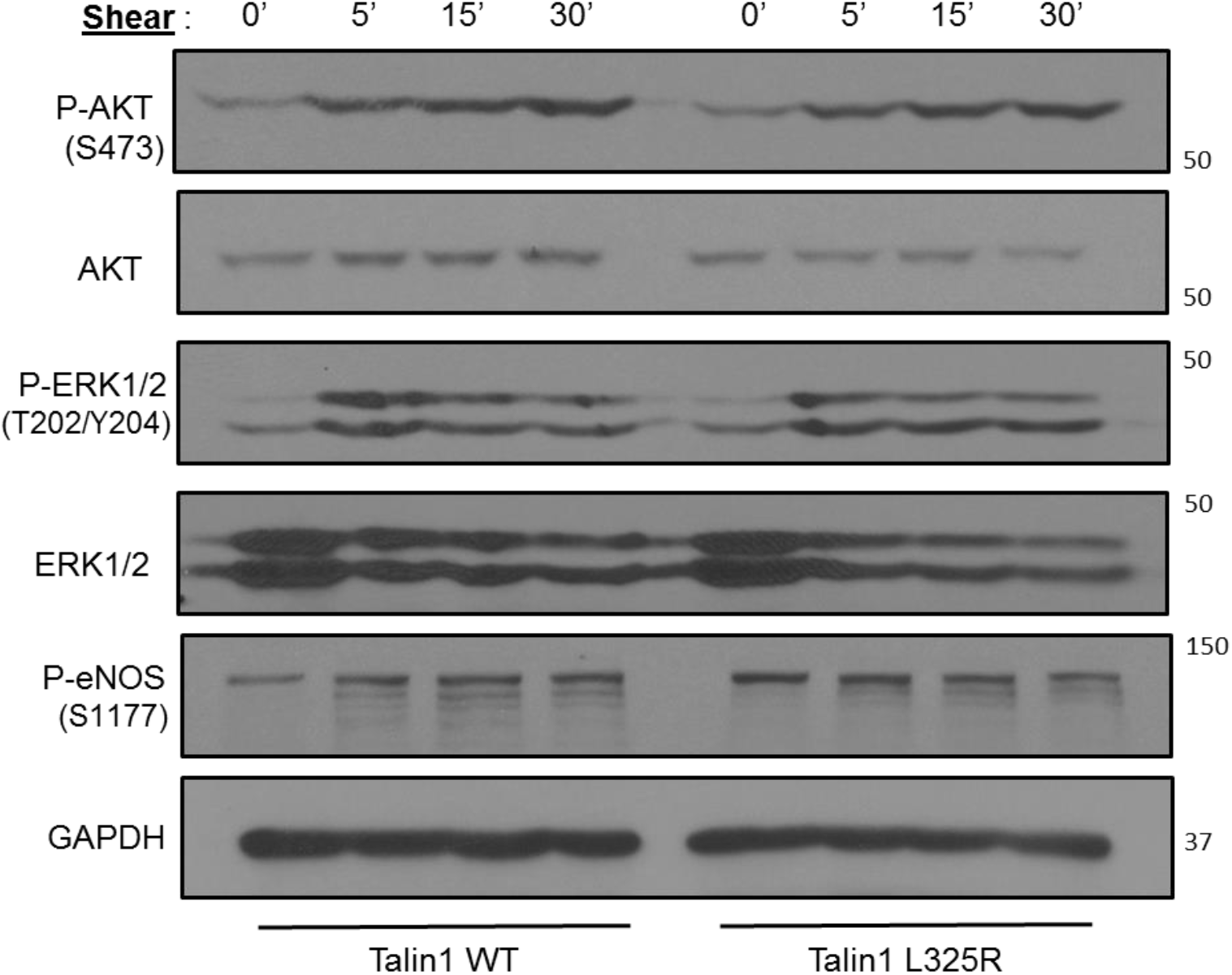
Talin1-dependent Integrin Activation isn’t required for Shear Stress-Induced AKT/ERK/eNOS signaling. Talin1 WT and Talin1 L325R cells were exposed to shear stress for the indicated time points. Western blot was done using phospho-specific antibodies to measure activation of ERK, AKT, and eNOS. Representative western blots are shown(n=3).

**Supplemental Figure VI.**
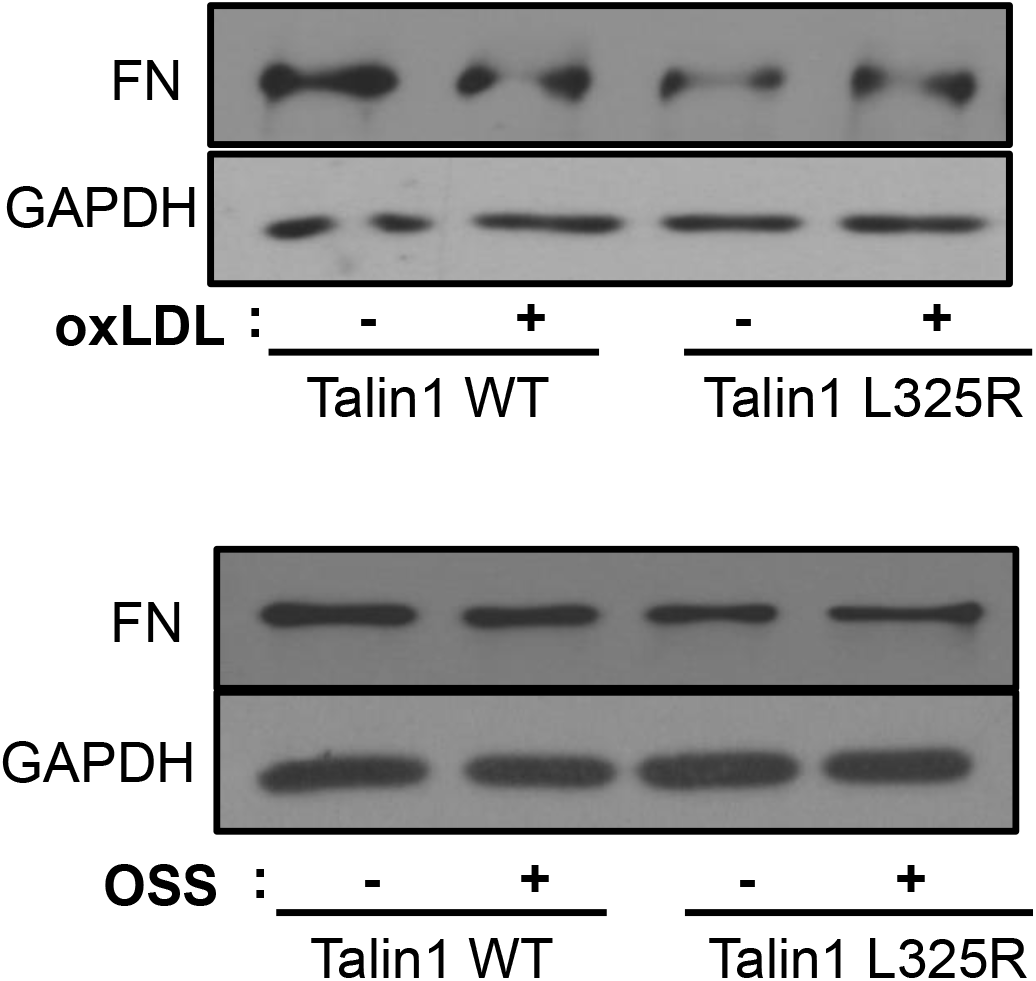
Intact Fibronectin Expression in Talin1 L325R Cells. Talin1 WT and Talin1 L325R cells were plated in Matrigel and then exposed to oxLDL or oscillatory shear stress for 24 hours.. Western blot was done using antibodies against fibronectin. n=3

**Supplemental Figure VII.**
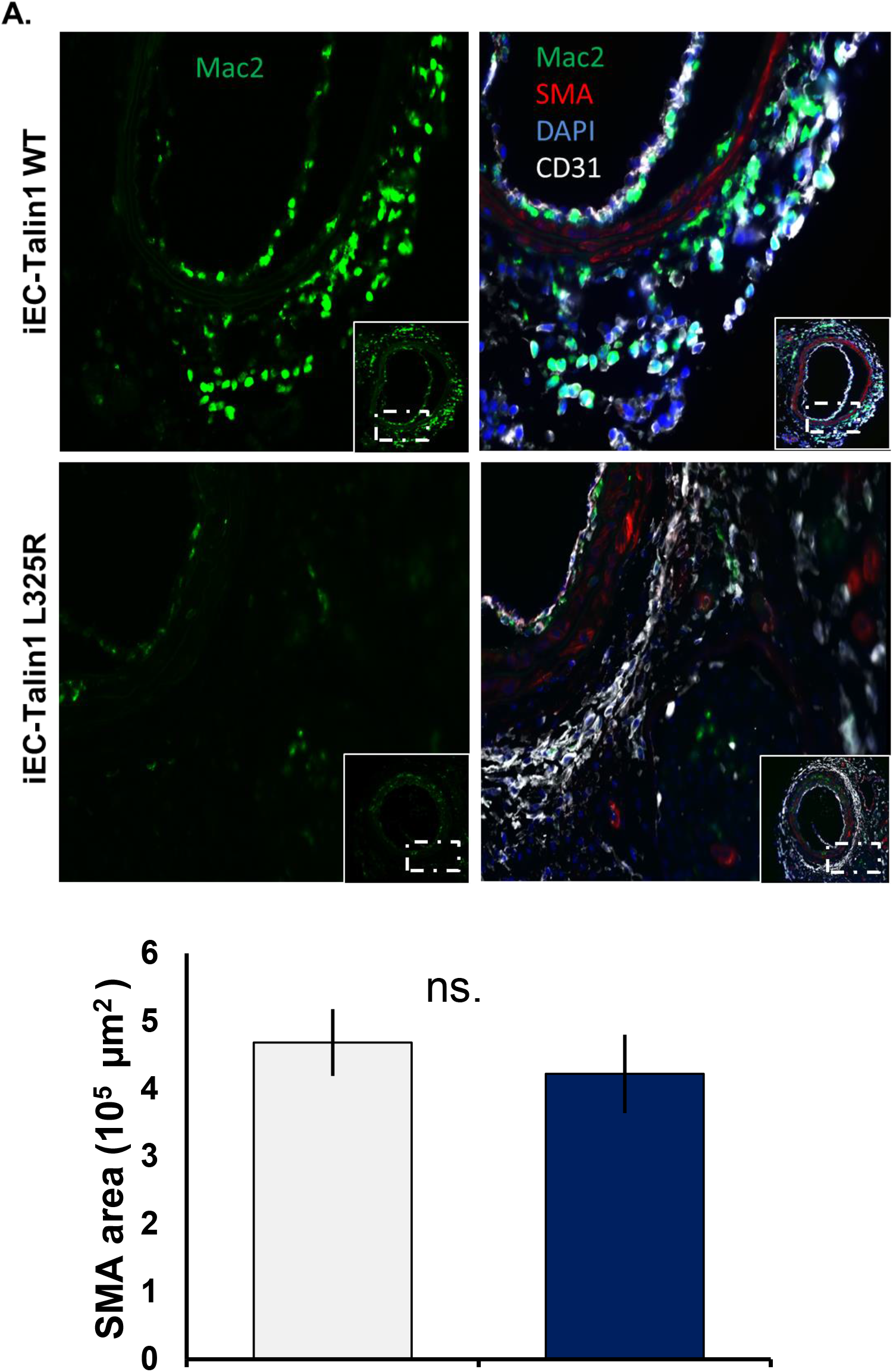
Impaired Talin1-Dependent Integrin Activation Does Not Affect Endothelial Coverage or Smooth Muscle Cell Content. After PCL for 7 days, immunohistochemistry was performed for MAC2(green) and SMA (red). Representative images are shown(n=5-6). Values are means ±SE. n.s. indicates not significant. Student t test was used for statistical analysis.

**Supplemental Table I.**
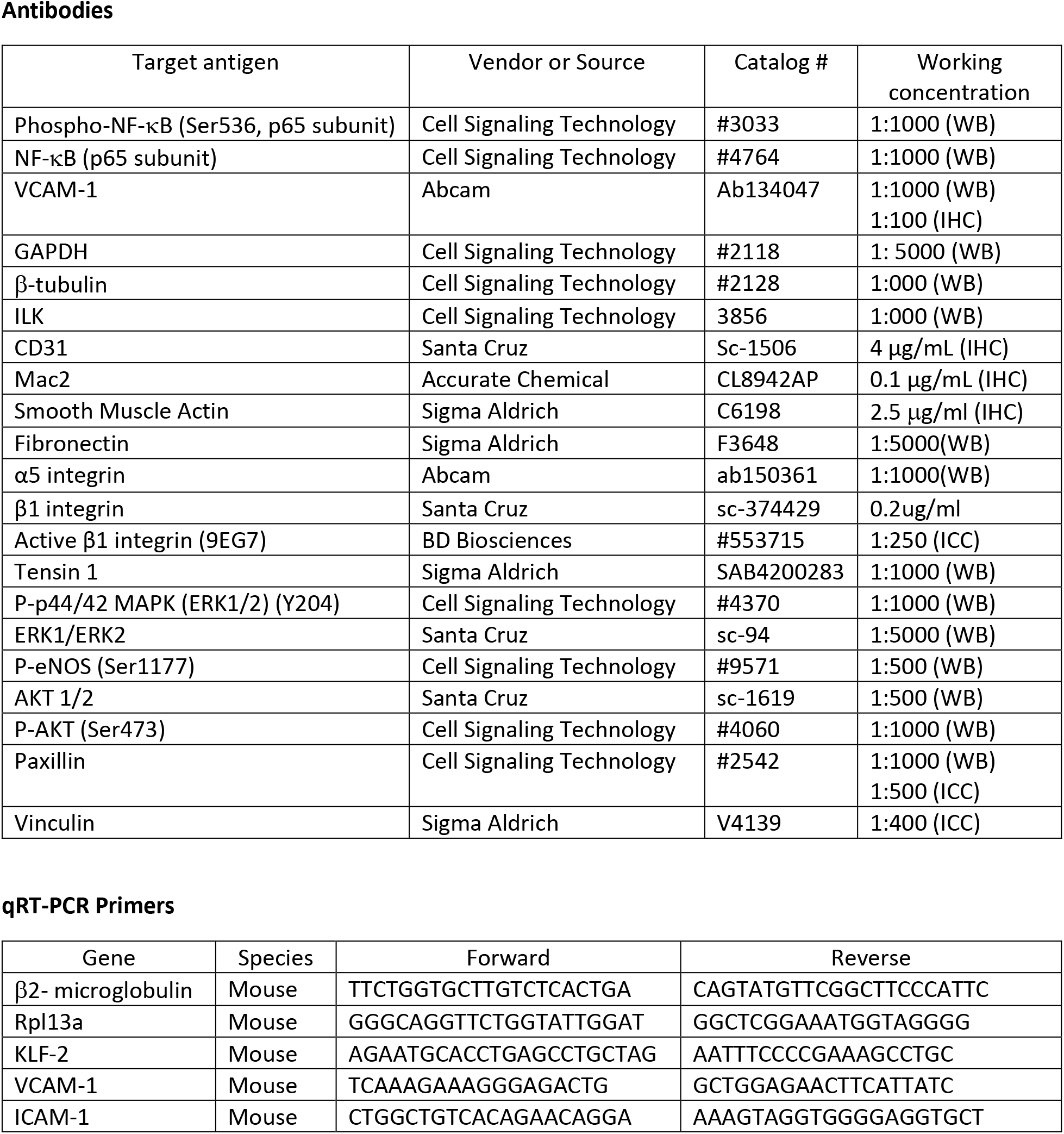

